# PUMA: PANDA Using MicroRNA Associations

**DOI:** 10.1101/2019.12.18.874065

**Authors:** Marieke L. Kuijjer, Maud Fagny, Alessandro Marin, John Quackenbush, Kimberly Glass

## Abstract

Conventional methods to analyze genomic data do not make use of the interplay between multiple factors, such as between microRNAs (miRNAs) and the mRNA transcripts they regulate, and thereby often fail to identify the cellular processes that are unique to specific tissues. We developed PUMA (PANDA Using MicroRNA Associations), a computational tool that uses message passing to integrate a prior network of miRNA target predictions with protein-protein interaction and target gene co-expression information to model genome-wide gene regulation by miRNAs. We applied PUMA to 38 tissues from the Genotype-Tissue Expression (GTEx) project, integrating RNA-Seq data with two different miRNA target predictions priors, built on predictions from TargetScan and miRanda, respectively. We found that while target predictions obtained from these two different resources are considerably different, PUMA captures similar tissue-specific miRNA-target gene regulatory interactions in the different network models. Furthermore, tissue-specific functions of miRNAs, which we identified by analyzing their regulatory profiles and which we made available through a Shiny app (https://kuijjer.shinyapps.io/puma_gtex/), are highly similar between networks modeled on the two target prediction resources. This indicates that PUMA consistently captures important tissue-specific regulatory processes of miRNAs. In addition, using PUMA we identified miRNAs regulating important tissue-specific processes that, when mutated, may result in disease development in the same tissue. PUMA is available in C++, MATLAB, and Python code on GitHub (https://github.com/kuijjerlab/PUMA and https://github.com/kuijjerlab/PyPuma).

## INTRODUCTION

The regulation of gene expression involves a complicated network of interacting elements. The biological process of transcription begins with the binding of transcription factors to specific sequence motifs upstream of a gene’s transcription initiation site. This induces conformational changes in the DNA and initiates the process of assembly of the RNA polymerase complex, which in turn carries out transcription of the gene to a messenger RNA (mRNA). At a post-transcriptional level, small non-coding RNA molecules such as miRNAs can repress mRNA translation and cause degradation of the mRNA transcript [1]. What emerges is not a single set of interactions, or even a single pathway, but a complex network of interacting genes and gene products. Capturing these interactions is critical as we seek to understand how gene expression is regulated in different tissue environments, and how this regulation is disrupted in disease.

MicroRNAs are small non-coding RNAs of about 22 base-pairs in length that can bind to the 3’ untranslated region (UTR) of their mRNA targets. The miRNA-mRNA duplex then associates with Argonaute family proteins, which recruit factors that induce mRNA degradation and translational repression [2, 3]. Most human protein coding genes are thought to be regulated by miRNAs, with over 60% having conserved miRNA binding sites in their 3’UTR [4]. miRNAs are generally thought to moderately downregulate their target genes, as individual sites usually reduce protein output by less than 50% [5]. However, most mRNAs have multiple miRNA regulatory sites in their UTR and miRNAs bound to these sites can act additively [6]. In addition, miRNAs may bind to many more non-canonical regulatory sites and act together, thereby increasing their regulatory potential [7].

Given the large number of miRNAs in the human genome (currently thought to be approximately 2,300 [8]) and because of their broad regulatory potential, their regulatory profiles are often modeled in gene regulatory networks. Such networks have generally been estimated using inverse correlation measures between expression levels of miRNAs and mRNAs. However, this approach has its limitations, as many different mechanisms modulate miRNA activity [9], and RNA transcripts may compete for binding to miRNAs, creating a more complex regulatory network than can be captured with co-expression patterns alone [10].

Other methods start with a prior network based on target predictions and then “color,” or assign weights to, the network’s nodes based on miRNA and mRNA expression levels [11]. However, target predictions are often different from actual interactions, with many studies reporting positive correlation between a miRNA and about half of its predicted targets [12]. Furthermore, target prediction remains challenging, and different algorithms may result in rather different networks of potential interactions [13]. Moreover, some genes may be regulated by a miRNA even though they are not predicted as a target of that miRNA by current prediction algorithms. Such “new” edges can not be learned if a model only considers known predicted targets.

Here, we present PUMA, or <u>P</u>ANDA <u>U</u>sing <u>M</u>icroRNA <u>A</u>ssociations, an algorithm that can directly model robust regulatory edges (including new edges) between miRNAs and their target genes, making use of prior knowledge on target predictions, fine-tuning these using information on co-regulation of the miRNA’s target genes. PUMA leverages the message passing framework described in our group’s previously developed algorithm PANDA [14], which models interactions between transcription factors and their target genes using message passing, thereby integrating multiple independent sources of data. This method starts with an initial estimate of the paths of information exchange between regulatory proteins (*i.e*. transcription factors) and their target genes. It then iteratively refines this prior network by incorporating gene expression and protein-protein interaction data, which provide information on the regulation of genes and on cooperative regulation by transcription factors, respectively. Since developing PANDA, we have used it to identify differences in transcriptional regulation between multiple human tissues [15], in transcriptonal regulation between tissues and their cells-of-origin [16], between ovarian cancer subtypes [17], and to identify sexual dimorphic gene regulation in colon cancer [18], among others [19–22].

While PUMA leverages the message passing framework used in PANDA, we introduced several critical modifications to incorporate the effects of miRNAs as an additional class of regulators into the gene regulatory network model, integrating miRNA target predictions alongside transcription factor regulatory predictions and gene expression levels using a modified message passing algorithm. We used PUMA to model miRNA regulatory networks for 38 tissues from the Genotype-Tissue Expression project (GTEx), integrating miRNA target predictions with gene expression data for each of the 38 tissues. We built two different collections of networks, each based on a prior obtained from a popular resource of miRNA target predictions, either TargetScan [23] or miRanda [24]. We extracted tissue-specific gene regulation by miRNAs, as well as miRNA functions from these two collections of networks. We found that PUMA consistently captures tissue-specific gene regulation by miRNAs, even when using different input sources of target predictions. Finally, we provide a new resource of tissue-specific functions of miRNAs identified with PUMA, and validate predicted tissue-specific functions in a database of disease-associated SNPs in miRNA target sites.

## MATERIALS AND METHODS

### The PUMA algorithm

We developed PUMA, a regulatory network reconstruction method to model miRNA-target gene interactions. PUMA models these interactions by integrating a regulatory prior with protein-protein interaction data and gene expression data. It uses an iterative message passing approach to model information flow between the different data types, finding “agreement” between data represented by multiple networks.

The method starts with a regulatory prior (*W*) of initial regulator-target gene interactions. These regulators can be either miRNAs or transcription factors (TFs). Initial regulatory interactions can be combined from multiple sources, such as miRNA-target predictions (for putative interactions of mRNAs by miRNAs), a TF motif scan (for estimated mRNA regulation by a TF), or ChIP-Seq data (for *in vivo* estimates of mRNA regulation by a TF). PUMA also uses a co-regulatory prior (*C*) of target gene co-expression levels measured using Pearson correlation on gene expression data, and has an optional input of initial TF-TF interactions, which can be based on known or predicted protein-protein interactions (PPIs). These PPIs are overlayed on an identity matrix that includes all regulators (*i.e.* all TFs and all miRNAs), resulting in co-operativity prior *P*.

PUMA performs a double z-score normalization on these three prior networks, and then quantifies the “agreement” between the different data types using a modified Tanimoto similarity score (*T*) [25], which evaluates the similarity between sets of interactions in two networks:

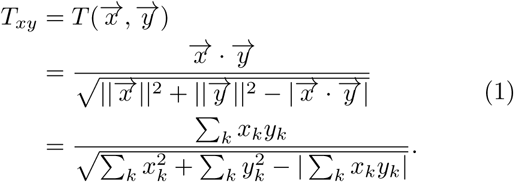

This score is used to calculate the “responsibility” (*R*) of an edge between a regulator *i* and a gene *j* at iteration step *t*. The responsibility estimate represents information flowing from a regulator to a target gene, and returns a confidence score for how strongly the target gene is regulated by this regulator, taking into account other potential regulators of the gene. PUMA uses protein-protein interaction information to estimate the cooperation between pairs of transcriptional regulators (TFs), and self-interactions to measure the responsibility by miRNAs, using the following equation:

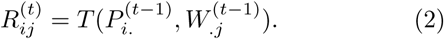

In a similar manner, the “availability” (*A*) estimate represents information flow from a target gene to a regulator and is based on the level of agreement between the targets of a regulator and the set of genes with which the target gene is co-regulated:

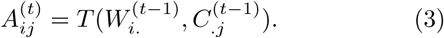

The initial regulatory network (*W*) is then updated using an update parameter *α* (with 0 < *α* < 1):

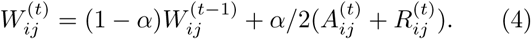

This is followed by updating the *P* and *C* networks. For each regulator (*i*), PUMA checks whether it matches with an entry in the input list of miRNAs (*q*). Since the only interactions a miRNA makes are through regulation of their targets (they do not act in complexes with other miRNAs or TF proteins), *P* will not be updated between any miRNA-TF interactions or between miRNA-miRNA interactions (except for self-interactions):

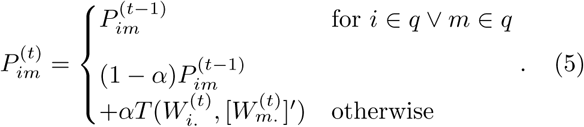

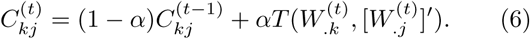

We note that self-interactions in *P* (including those among miRNAs) and *C* are then separately updated in order for the algorithm to converge, as in Glass *et al.* [14, 26]. These message passing steps are repeated until the regulatory network converges.

The message passing framework presented here and used by PUMA, is highly similar to the one from the PANDA algorithm [14, 26]. However, it includes several critical adjustments to account for the mechanisms of miRNA regulation. In particular, we have modified both the initial co-operativity network and its update (as we show in Equation (5)) to account for the different types of regulatory behaviors of TFs and miRNAs. For more details on the message passing algorithm we implemented in PUMA, please refer to Glass *et al.* [14, 26].

PUMA is available in C++ and MATLAB code at https://github.com/kuijjerlab/PUMA, and as Python code at https://github.com/kuijjerlab/PyPuma.

### GTEx RNA-Seq data

We downloaded the Genotype-Tissue Expression (GTEx) version 6.0 RNA-Seq data (phs000424.v6.p1, 2015-10-05 released) from dbGaP (approved protocol #9112). GTEx release version 6.0 sampled 551 donors with phenotypic information and included 9,590 RNASeq assays (Consortium, 2015). We used our previously described method YARN [27] to perform quality control, which removed samples with sex-misidentification and merged related sub-tissues, resulting in a dataset of 9,435 gene expression profiles in 38 tissues from 549 individuals.

We used default settings in YARN to perform gene filtering and tissue-aware normalization using qsmooth [28]. However, only 85 pre-miRNA transcripts were retained after the filtering step in YARN. Because we were particularly interested in using miRNA expression levels to assess the properties of tissue-specific regulator miRNAs that we identified using our networks, we repeated the YARN pipeline without filtering out miRNAs. This resulted in normalized expression levels for 31,384 transcripts, which included 1,136 miRNAs.

To ensure that this procedure did not significantly alter the expression levels obtained with the standard YARN pipeline, we compared expression levels of the 30,248 mRNA and the 85 pre-miRNA transcripts that were not filtered by the standard YARN pipeline with their values in the dataset in which we included all pre-miRNAs. mRNA transcripts correlated with median Pearson *R* = 0.9982, range [0.9677, 1] and the 85 pre-miRNAs correlated with median Pearson *R* = 0.9999, range [0.9897, 1], indicating that adding counts of pre-miRNAs to the count data before the normalization step did not significantly alter the normalized expression levels of other genes.

### Pre-processing miRNA target prediction data

We downloaded miRNA target predictions from TargetScan v7.1 (all predictions, file “Summary Counts.all predictions.txt.zip,” http://www.targetscan.org/cgi-bin/targetscan/data_download.vert71.cgi, accessed: July 8, 2016) and miRanda (predictions with “Good mirSVR score, Conserved miRNA,” file “human predictions S C aug2010.txt,” http://34.236.212.39/microrna/getDownloads.do, accessed: December 5, 2017). We filtered TargetScan interactions by selecting Homo sapiens interactions and by removing interactions with context+ scores larger than −0.1, resulting in 1,524 miRNAs and 18,234 target genes. The miRanda prior contained 1,100 miRNAs and 19,796 target genes.

We selected miRNAs that were present as regulators in both the TargetScan and miRanda priors, and for which expression levels were available. To do this, we first needed to match miRNA identifiers between the different types. We converted miRNA regulator identifiers from TargetScan and miRanda to gene names by changing the character vector to uppercase, removing the “hsa-”prefix, removing the dash character after “LET” and “MIR,” and pasting “MIR” in front of miRNAs that start with “LET.” We removed all extensions to obtain a list of “base” miRNAs, of which 578 were present as regulators in both prior target prediction resources, and were also available in the expression data.

The numbered suffix in the miRNA identifiers indicate diverse loci that produce identical mature miRNAs. We therefore collapsed these miRNAs in the prior by taking the union of all edges. For duplicates, we selected the most significant edges (lowest context+ score for TargetScan, all edges for miRanda). 643 “regulator” miRNAs corresponded to the set of 578 “base” miRNAs.

An asterisk (*) extension indicates an alternative transcript with lower expression levels. However, information on expression levels of mature miRNAs could have been derived from experiments in specific cell lines or under specific experimental settings, and their expression levels may vary in different tissues. miRNAs with asterisk extensions were only present in miRanda, not in TargetScan. To be able to match these miRNAs between the two different priors, we merged those in the miRanda prior by taking the union of edges.

The −3p/−5p extension in miRNA identifiers indicates whether the mature miRNA product comes from the 3’ or 5’ end of the hairpin structure formed by the pre-miRNA [3]. This indicates a different mature product with a different seed sequence (that may target different genes). These miRNAs are supposed to have similar expression levels [29], although recent reports identified imbalance in −3p/−5p expression ratios [30]. When evaluating the expression levels of such miRNAs, we used the expression level associated with the miRNA’s gene name, so that the same expression level values were assigned to miRNAs from the same genomic location.

This preprocessing resulted in a set of 621 “target” miRNAs for which we had expression data available, which corresponded to the set of 578 “base” miRNAs. Finally, we took the intersection of the lists of target genes in the TargetScan and miRanda regulatory priors with genes for which we had expression data available, resulting in 16,161 target genes.

### Regulatory network reconstruction

We used the MATLAB version of PUMA to integrate target predictions from TargetScan and miRanda with gene expression data from each of the 38 GTEx tissues to estimate gene regulatory networks for each of these tissues. In total, we modeled 76 gene regulatory networks, two for each tissue.

### Comparison of tissue-specific edges

PUMA returns complete, bipartite networks with edge weights similar to z-scores. To compare the tissue-specificity of network edges, we calculated a tissue-specific edge score, which was defined as the deviation of an edge weight 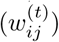 between a miRNA (*i*) and a target gene (*j*) in a particular tissue (*t*) from the median of its weight across all tissues, using the interquartile range (IQR) (as in Sonawane *et al.* [15]):

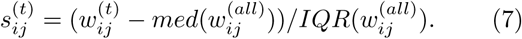

We defined an edge with a specificity score 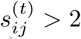 as specific to tissue *t* and the multiplicity of an edge as the number of tissues it is specific to:

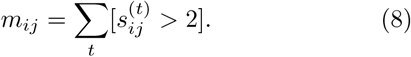

To determine tissue-specific expression levels of miRNAs, we compared the median expression level 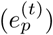 of a miRNA (*p*) in a particular tissue (*t*) to the median and IQR of its expression levels across all tissues (similar to what we described in Sonawane *et al.* [15]):

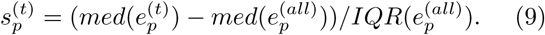

We assessed the overlap of the initial TargetScan and miRanda gene regulatory priors with the Jaccard index and with Pearson correlation. We compared tissue-specific edge scores from networks reconstructed on the two different priors using Pearson correlation.

### Gene set enrichment analysis on miRNA targeting profiles

For each miRNA in a given tissue, we selected its tissue-specific targeting profile, meaning all tissue-specific scores connected to that miRNA. We ran a pre-ranked Gene Set Enrichment Analyses (GSEA) [31] on these profiles to test whether miRNAs specifically target Gene Ontology (GO) terms in the different tissues. We ran GSEA for networks reconstructed on the TargetScan prior, as well as for networks reconstructed on the miRanda prior. Thus, in total, we ran 48,868 GSEA analyses.

For each miRNA/tissue pair, we calculated tissue-specific GO term enrichment scores, which we defined as the −*log*_10_*FDR* (False Discovery Rate) from GSEA, multiplied by the sign of the GSEA Enrichment Score (ES)—those with *ES* < 0 were multiplied by −1, those with *ES* > 0 were multiplied by 1. We then used Pearson correlation across all 733 tissue-specific GO term enrichment scores for each miRNA/tissue pair to assess the similarity of tissue-specific regulation of biological processes by miRNAs.

### Community structure analysis to identify sets of related tissue/miRNA GO terms

We selected highly significant (FDR *<* 0.001) and positively enriched (Enrichment Score *>* 0.65) associations from these analyses and converted these scores into a binary matrix. We then used fast-greedy community detection [32] on this matrix to cluster the data and to identify communities or network modules that share tissue-specific regulatory patterns.

We then used the Jaccard index to compare nodes (miRNA/tissues and GO terms) that belonged to communities that included at least 5 GO terms in either the TargetScan or the miRanda networks. We used word clouds to visualize the tissue-specific functions of miRNAs in these communities. To do this, we split the strings for each of the significant GO term into separate words and removed words that occurred less than 3 times to obtain a background list of word frequencies associated with all significant GO terms. We then counted the number of times a word was present in the community of interest, and divided this by the total number of words associated with significant GO terms in that community (the “observed” rate), as well as the number of times the word occurred in the background list, divided by the total number of words in that background list (the “expected” rate). We then calculated the observed/expected ratio, multiplied this by 10, and rounded this number to an integer to obtain a word occurrence score. Finally, we added the word, repeating it by its word occurrence score, to a list. We used this list of normalized word occurrences as input in https://www.wordclouds.com/ to generate a word cloud for that community. We repeated this for each of the communities that included tissue-specific targeting by miRNAs of at least 5 GO terms.

### Data availability

The reconstructed networks are available on Zenodo (doi: 10.5281/zenodo.1313768; https://tinyurl.com/puma-gtex). An R Shiny app [33] that can be used to assess tissue-specific functions of miRNAs using different thresholds is hosted on https://kuijjer.shinyapps.io/puma_gtex/.

## RESULTS

### Tissue-specific gene regulation by miRNAs

We started by reconstructing miRNA-target gene genome-wide regulatory networks for a large collection of human tissues. We downloaded RNA-Seq data for 54 different tissues (including three different cell types) using Bioconductor package YARN [27]. Within YARN, we performed quality control and normalization of the data, merging tissues with similar expression profiles (see Methods), which retained gene expression data for 9,435 samples across 38 tissues. We limited network reconstruction to only those genes and miRNAs that were expressed and which appeared in the TargetScan and miRanda prior, leaving 16,161 genes and 621 miRNAs; these 621 target miRNAs corresponded to 643 regulators in the prior networks (see Methods). We then used PUMA to integrate target gene co-expression information for each tissue with an initial regulatory network, which we based on either TargetScan or miRanda miRNA target predictions. Consequently, our analysis provides two alternative miRNA mediated gene regulatory networks for each of the 38 tissues (tissue-networks), one based on the TargetScan prior and the other alternative based on the miRanda prior.

We tested for tissue-specific edges in these networks. We defined an edge to be tissue-specific if its weight was larger than twice the interquartile range of its weight across all 38 networks (see Methods). We identified highly similar numbers of tissue-specific miRNA-target gene regulatory edges in the networks modeled on the two different priors—3.093 million and 3.098 million edges for networks modeled on the TargetScan and miRanda prior, respectively (see Figure 1). In addition, the proportion of tissue-specific edges identified in the different tissues was comparable between the networks modeled on the two different priors (Pearson *R* = 0.92). The proportion of multiplicities, or the number of tissues in which an edge is identified as specific, was also similar between the two different models.

**Figure 1.**
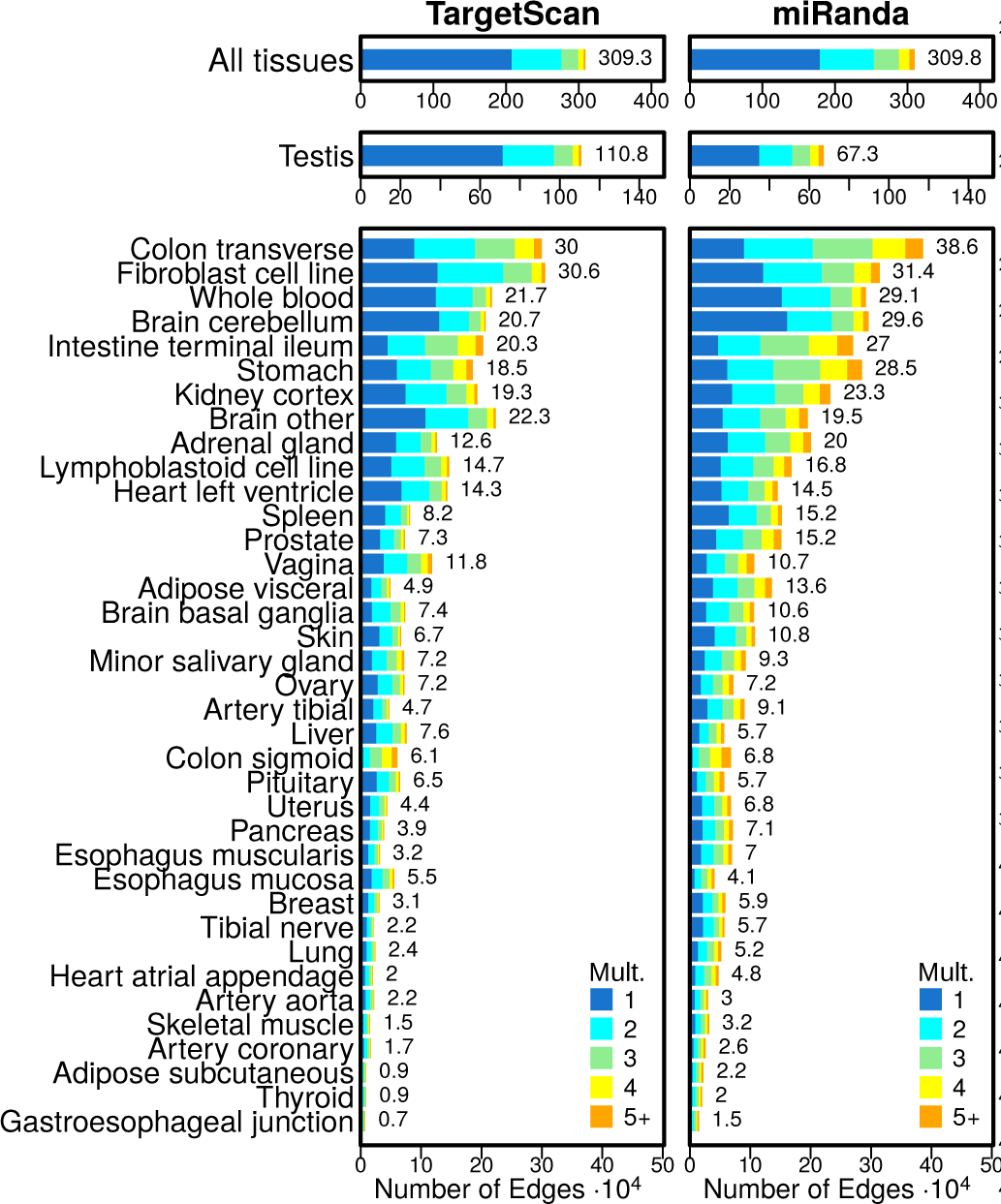
Bar plots illustrating the number of edges modeled on the TargetScan and miRanda priors. The number of elements identified as specific in each tissue is shown to the right of each bar. Tissues are ordered by the average number of tissue-specific edges. Mult.: the edge multiplicity, or the total number of tissues an edge is specific to.

Even though we identified approximately the same total number of tissue-specific edges across all tissues, on average we identified a larger number of tissue-specific edges per tissue in the networks modeled on the miRanda prior (t-statistic= 7.1, p-value= 6.5 · 10^−10^). However, we found the opposite to be true for testis, the tissue with the greatest number of tissue-specific edges in both priors. In testis, we identified a substantially larger number (1.65×) of tissue-specific edges in the networks modeled on the TargetScan prior than in the networks modeled on the miRanda prior.

We next assessed how similar the edge tissue-specificity scores were between the networks modeled on the different priors. For each of the 38 tissues, we calculated the Pearson correlation coefficient on edge tissue-specificity scores between the tissue-network modeled on the two different priors. We found that, in general, PUMA networks modeled using different target predictions result in similar tissue-specificity scores (median Pearson *R* = 0.63, Figure 2A and Supplemental Figure S1). For all tissues, except testis, the resulting PUMA tissue-networks modeled on the two different priors were more similar than the two prior networks (*R* = 0.34). This means that, even though there may be differences between various target prediction resources, PUMA helps fine-tuning these predictions into tissue-specific regulatory interactions. We believe that the anomalous gene expression patterns observed in testis, which has been described previously [15], may, at least in part, be caused by differential targeting by miRNAs.

**Figure 2.**
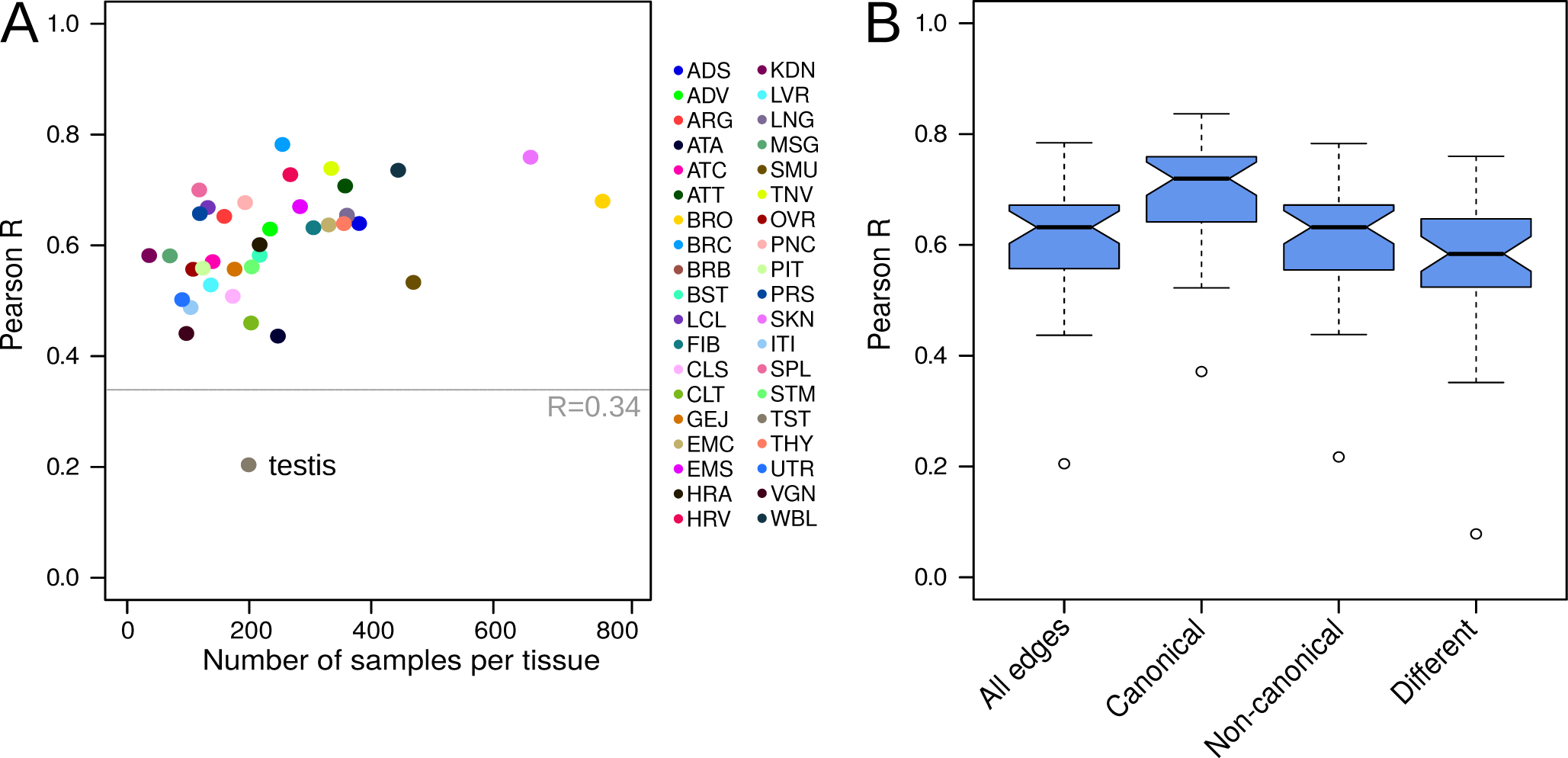
A) Tissue-specificity score similarity—measured using Pearson correlation coefficient (Pearson R)—for each of the 38 miRNA gene regulatory tissue-networks modeled on TargetScan and miRanda priors, compared to the number of samples available for each tissue. TargetScan and miRanda priors correlate with *R* = 0.34. B) Boxplots depicting the distribution of edge similarity for all edges, edges that are canonical in both priors, edges that are non-canonical in both priors, and edges that are different between the TargetScan and miRanda priors. Boxplots represent the median and IQR, with whiskers extending out from the box to 1.5× the IQR.

We then examined the similarity between tissue-specificity scores for miRNA-target gene interactions that were predicted by both TargetScan and miRanda (“canonical” interactions), interactions that were neither predicted in TargetScan nor in miRanda (“non-canonical” interactions), and edges that were predicted interactions in one of the priors but not in the other (“different,” or inconsistent interactions). We used Pearson correlation to evaluate the similarity of these different types of edges. We found that, in general, tissue-specificity levels of edges that were canonical in both priors were most reproducible, followed by edges that were non-canonical in both priors. As expected, miRNA-target gene interactions that were canonical in only one of the two priors were less similar than edges that were canonical or non-canonical in both priors. However, for those edges the median similarity was still *R* = 0.58 (Figure 2B). This indicates that PUMA can capture consistencies in miRNA-target gene regulation, even when there are inconsistencies between different target prediction resources, and highlights the strengths of modeling miRNA target gene interactions with PUMA.

### Tissue-specific miRNA targeting patterns

To better understand tissue-specific functions of miRNAs, we ran pre-ranked gene set enrichment analysis on the tissue-specific targeting profile of each miRNA in each of the 38 tissues. We did this both for the collection of tissue-networks modeled using the TargetScan and for networks modeled using the miRanda priors (see Methods). We calculated the tissue-specific targeting scores of all 733 available GO terms, and investigated whether tissue-specific regulation of biological processes by miRNAs was similar in the two different collections of networks. Most (89.6%) of the miRNA/tissue pairs had a positive Pearson correlation coefficient, with a median Pearson R of 0.66 (see Figure 3 for the correlations between all GSEA scores, and Supplemental Figure S2 for the correlations separated by tissue).

**Figure 3.**
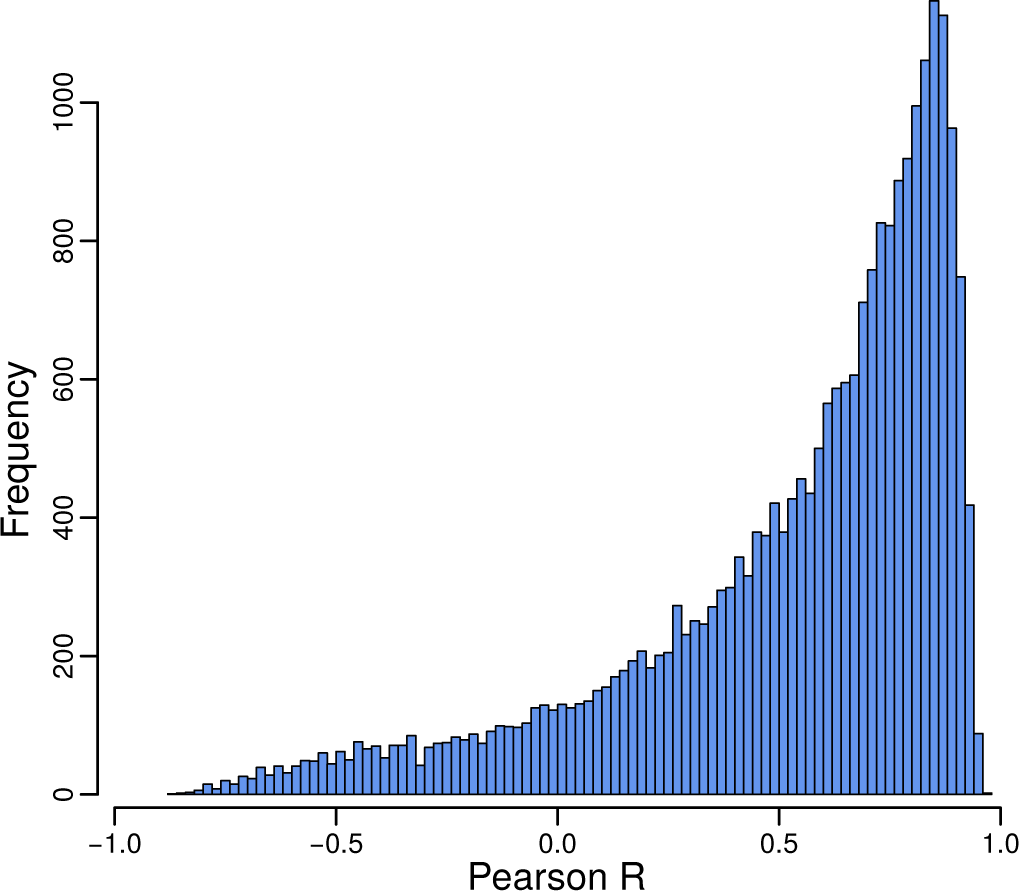
Pearson correlation between the GSEA scores obtained from the 24,434 tissue-specific miRNA targeting profiles computed on TargetScan and the 24,434 profiles computed on the miRanda prior. The bulk of miRNAs have correlating GO term scores in networks modeled using different miRNA priors.

As a negative control, we computed the correlation of miRNA–GO term GSEA scores between different tissues, for the miRanda and TargetScan-generated networks separately. The resulting correlation coefficients were centered around zero, with median Pearson *R* = −0.002 for the miRanda networks and *R* = −0.001 for the TargetScan networks, respectively (Supplemental Figure S3). GSEA scores for miRNA/tissue pairs obtained from the two different collections of networks were significantly more correlated than the negative control (2-group Wilcoxon signed-rank test p-value < 2.2 · 10^−16^). These results confirm that, even though we used different target predictions as input for PUMA, the actual tissue-specific regulatory functions we obtain from analyzing these networks are highly similar.

We tested whether similar miRNA/tissue pairs, as identified in both models, control similar biological functions. To do this, we selected highly significant miRNA/tissue–GO term associations (*FDR* < 0.001, *ES* > 0.65), and performed community structure analyses on these sets of associations to identify shared tissue-specific targeting patterns of miRNAs across the tissues. We identified 67 communities (*e.g.* sets of GO terms grouped together with miRNA/tissue pairs) in the regulatory associations identified in PUMA networks modeled on the TargetScan prior, and 64 communities in those identified in PUMA networks modeled on the miRanda prior. The overall modularity of these community structures was 0.76 and 0.77, respectively (see Figure 4A–B).

**Figure 4.**
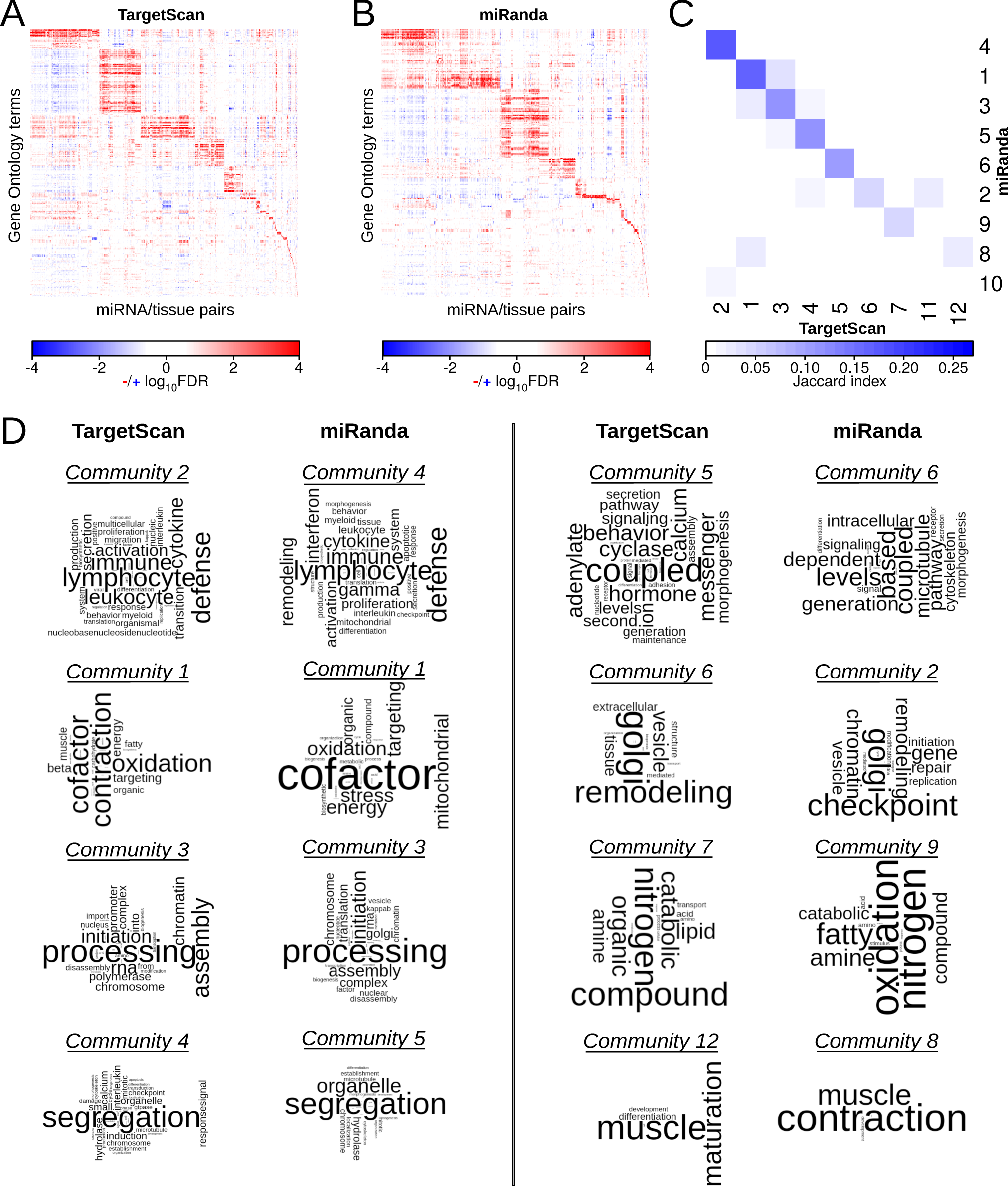
A–B) Heatmaps depicting communities of significantly targeted GO terms (*FDR* < 0.001, *ES* > 0.65) based on GSEA analyses on all possible miRNA/tissue pairs for the networks modeled on the TargetScan (A) and the miRanda (B) prior. C) Similarity (measured with Jaccard index) of miRNA/tissue–GO term associations in communities targeting at least five GO terms identified in networks modeled using the TargetScan or miRanda prior. D) Word clouds depicting communities targeting at least five GO terms. Community pairs with the highest Jaccard index are shown. We omitted TargetScan community 11 and miRanda community 10 as they each mapped to communities that corresponded to another community (6 and 4, respectively) with a higher Jaccard index.

In both collections of PUMA tissue-networks, nine communities were associated with at least five GO terms. For each of these communities, we calculated the Jaccard index between the two different sets of miRNA/tissue– GO term associations to evaluate the overlap in miRNA/tissue pairs and GO terms associated with the community (Figure 4C). As can be seen from this figure, the communities have relatively high node overlap, indicating that similar processes are identified as regulated in a tissue-specific manner by similar sets of miRNAs in both analyses. We used word clouds to visualize the biological processes that were associated with these communities, which allowed us to further explore these similarities. Figure 4D shows that similar biological processes are identified as regulated by miRNAs in a tissue-specific manner in these communities. These include processes involved in the immune system, mitochondrial respiration, translation initiation, chromosome segregation, intracellular signaling, protein transport, and muscle contraction.

Importantly, we can identify these communities of similarly regulated biological processes in networks modeled using different prior target predictions. This indicates that PUMA’s message passing allows us to discover patterns of tissue-specific regulation by miRNAs, even though there may be inconsistencies in the initial target predictions that we used as prior input in PUMA.

### A resource of tissue-specific miRNA functions

We compiled a resource of miRNAs that regulate biological processes in a tissue-specific manner. To do this, we took the union of miRNA/tissues significantly regulating GO terms in the TargetScan and the miRanda networks (8,992 miRNA/tissue–GO terms in total). We then subsetted this list to only those miRNA/tissues for which the tissue-specific targeting profiles correlated with *R* > 0.8 (2085 miRNA/tissue–GO terms). This list of significant tissue-specific functions of miRNAs contained 423 regulator miRNAs, 37 tissues (no consistent tissue-specific regulation was identified for tibial nerve), and 174 GO terms. This resource can be accessed at https://kuijjer.shinyapps.io/puma_gtex/.

We assessed over-representation of miRNAs, GO terms, and tissues in this resource of significant interactions (Figure 5A–C). Twenty-one miRNAs were over-represented (> *median* + 2 *IQR*) in regulating multiple tissue-specific processes (Figure 5A). MIR517C and MIR1468 were the two miRNAs with the highest number of associations of tissue-specific regulation of biological processes (Figure 5A). MIR517C was associated with immune system processes in artery tibial and thyroid, with “regulation of neurogenesis” and “regulated secretory pathway” in “brain other” (a compound of multiple brain regions), with synapse-associated processes and “extracellular structure organization and biogenesis” in skeletal muscle, with sperm-associated pathways in testis, and with “ovulation cycle” and ectoderm-associated pathways in vagina (Figure 5D–E). This miRNA has been detected in maternal plasma. It was also recently described to be overexpressed in parathyroid carcinoma [34] and to inhibit autophagy and epithelial-to-mesenchymal transition in glioblastoma, a malignant brain cancer [35], indicating that deregulation of the expression of this miRNA in tissues in which it regulates tissue-specific processes may lead to cancer.

**Figure 5.**
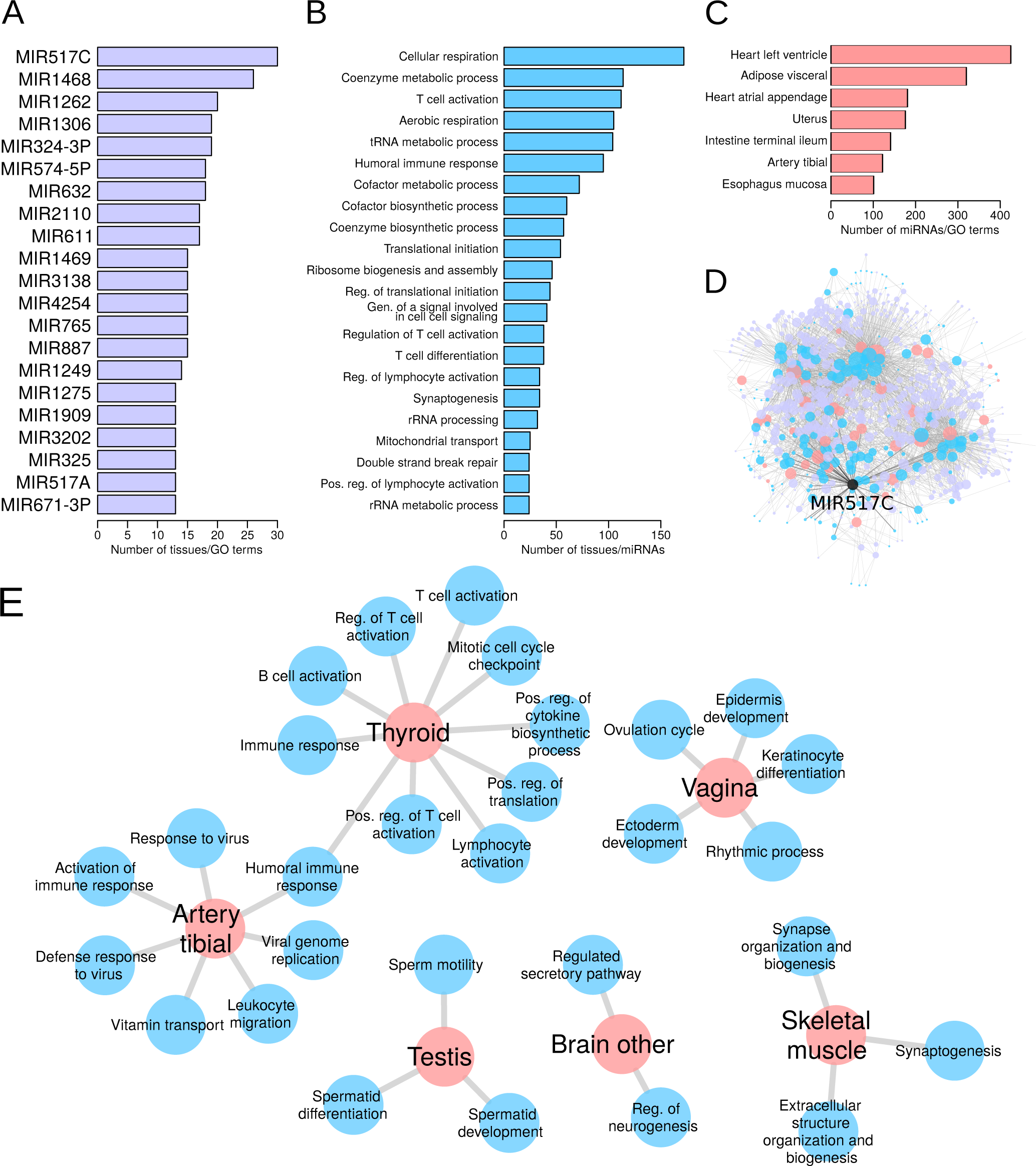
Over-represented miRNAs (A), GO terms (B), and tissues (C) in the database of significant tissue-specific functions of miRNAs. D) Visual representation of the significant tissue-specific regulatory functions of miRNAs present in the Shiny database. Node color illustrates tissues (peach), GO terms (cyan), and miRNAs (magenta). Node size corresponds to the node’s degree. MIR517C—the miRNA with the largest over-representation in tissue-specific connections—is highlighted in black. E) Significant tissue-specific connections made by MIR517C. Reg.: regulation; Gen.: generation; Pos.: positive.

MIR1468 was associated with “sperm motility” in testis, with many immune system processes in thyroid, and with “double strand break repair” and chromatin-associated processes in whole blood. This miRNA has been implicated in different cancer types [36, 37] and the latter pathway may indicate a potential mechanism for this. In fact, MIR1468 was recently shown to promote tumor progression by activating PPAR-*γ*-mediated Akt signaling in hepatocellular carcinoma [38].

Twenty-two processes were more often targeted in a tissue-specific manner by miRNAs in more tissues than expected by chance. Most of these processes play a role in respiration and metabolism, immune response, and protein translation (Figure 5B), indicating that miRNAs play an important role in regulating these pathways in a tissue-specific manner in multiple tissues.

Seven tissues received significantly more tissue-specific gene regulation by miRNAs compared to all tissues (Figure 5C). Tissues receiving most tissue-specific gene regulation by miRNAs include heart left ventricle, adipose visceral, and heart atrial appendage. We do not know why these tissues have a higher amount of tissue-specific gene regulation by miRNAs. It may be that these tissues are more highly differentiated than others because of the specialized functions they carry out, and so the elevated miRNA activity represses extraneous functions. This could be a potential new area for research.

### miRNAs regulating tissue-specific processes are not differentially expressed

We wanted to evaluate whether tissue-specific regulation by miRNAs was caused by tissue-specific expression of those miRNAs. We identified 423 (66%) miRNAs that regulate biological processes in a tissue-specific manner. These regulator miRNAs were associated with 309 different miRNA genes. We compared the expression levels of these 309 miRNAs with those of the remaining 312 miRNAs, and found that miRNAs regulating biological processes in a tissue-specific manner have overall higher expression levels across all samples (two-sided Wilcoxon rank sum test statistic = 4.35 10^12^, p-value=2.2·10^−16^).

However, when comparing the tissue-specificity scores of these miRNAs in the tissue in which they regulate biological processes, we did not identify any association. The mean tissue-specificity score (difference in median expression in tissue-of-interest compared to overall median expression, divided by IQR) of these miRNAs was 0.022 (range [−0.820, 6.811]), indicating that these miRNAs were not specifically expressed in the tissue they regulate. While none of the miRNAs met our threshold of tissue-specific underexpression, six miRNAs had tissue-specificity scores larger than 2, suggesting tissue-specific overexpression of these miRNAs. These included MIR142-3P regulating the “insulin receptor signaling pathway” in spleen, MIR1909 regulating “coenzyme biosynthetic process” in testis, MIR200B regulating “aerobic respiration” and “cellular respiration” in pancreas and “regulation of muscle contraction” in prostate, MIR203 regulating “translational initiation” in esophagus mucosa, MIR208A regulating “activation of immune response”, “adaptive immune response”, “humoral immune response”, and “synaptogenesis” in heart atrial appendage, and MIR632 regulating “spermatid differentiation” in testis.

These findings are in line with our previous results in transcriptional regulatory networks, in which we identified no clear association between a transcription factor’s expression level and its tissue-specific regulation of biological processes [15]. They are also consistent with our previous finding that modeling of transcriptional gene regulatory networks is able to identify biologically relevant differences in regulatory processes even in situations where there is little or no differential expression [18]. Importantly, our results indicate that analysis of miRNA-mRNA co-expression networks, while potentially informative in identifying co-regulation of miRNA and mRNA expression levels, may miss miRNAs that are not differentially expressed, but that do regulate their targets in a tissue- or disease-specific manner.

### miRNA functions correspond to disease-associated SNPs

To further validate our findings, we integrated the tissue-specificity scores with miRdSNP, a database of Single Nucleotide Polymorphisms (SNPs) in the 3’UTR of human genes [39]. To do this, we downloaded the miRdSNP database, converted and matched miRNA names, and intersected miRNAs and target genes present in miRdSNP with those present in our regulatory networks. We then matched diseases listed in miRdSNP to GTEx tissues (manual curation). This left us with 24 GTEx tissues for which miRNA-target gene associations with disease were available. For each of these 24 tissues, we took the edge with the highest tissue-specificity score based on the TargetScan networks (range [−0.76, 2.68], mean= 1.11) and obtained the miRNA’s tissue-specific functions from our Shiny app (settings −*log*_10_(*FDR*) < 0.8). This resulted in tissue-specific pathway associations for 18/24 miRNAs. We then compared the pathways with the highest enrichment (based on ES) with the target gene’s description in GeneCards (www.genecards.org).

We found that the tissue-specific function of miRNAs obtained from the PUMA networks often matched that of the gene that had disease-associated SNPs in their miRNAs binding sites. For example, the pathway with the highest level of tissue-specific regulation by MIR429 in coronary artery was “endothelial cell migration.” The gene associated with the disease edge from miRdSNP was *VEGFA*, which encodes for a receptor important in angiogenesis. MIR140-5p specifically regulates “acute inflammatory response” in sigmoid colon. This miRNA was associated with disease SNPs in *TLR4*, an receptor involved in innate immunity. In ovary, MIR429 specifically targets “G1 phase of mitotic cell cycle” and was associated to SNPs in *CDK2*, a cell division gene. MIR1197 specifically targets “icosanoid metabolic process” in pancreas, and had disease-associated SNPs in *ADIPOR2*, a gene involved in glucose and lipid metabolism. All disease-associated edges are listed in Supplemental Table S1.

These results indicate that the tissue-specific functions of miRNAs predicted using PUMA are important for maintaining tissue homeostasis, and that disrupting miRNA-target gene edges in the regulatory network can perturb these processes, thereby influencing disease. This highlights the importance of modeling genome-wide miRNA-target gene regulatory networks in human tissues.

## DISCUSSION

In this manuscript, we describe PUMA, a new method to model gene regulation by miRNAs. PUMA integrates target gene co-expression information with initial target predictions, which can be obtained from resources such as Targetcan or miRanda. We applied PUMA to a large-scale RNA-Seq dataset from GTEx to identify tissue-specific regulatory patterns of miRNAs. We modeled two different collections of tissue-networks by integrating gene expression data from GTEx with two prior datasets—target predictions from TargetScan and miRanda, two of the most widely used miRNA target prediction resources. We found that tissue-specific gene regulation by miRNAs was reproducible for most tissues, except for testis. Potentially, the aberrant gene expression pattern in testis is, at least in part, caused by differential regulation by miRNAs. While tissue-specificity of gene regulation was reproducible for different types of edges, it was highest for edges that were predicted in both priors, indicating that compendium-like approaches using the intersection of different miRNA target prediction resources as prior data for network modeling could result in more accurate results.

We performed high-throughput gene set enrichment analyses on the tissue-specific targeting profiles of each of the miRNAs to characterize tissue-specific regulation of biological processes. We found that tissue-specific regulation of biological processes by miRNAs was highly similar in the networks modeled on different priors. We highlighted biological processes that were regulated in a tissue-specific manner (by different sets of miRNAs) in multiple tissues (Figure 4). The processes we identified play a role in the immune system, mitochondrial respiration, translation initiation, chromosome segregation, intracellular signaling, protein transport, and muscle contraction. In addition, we identified miRNAs and tissues for which we found an over-representation of tissue-specific regulation. The most enriched tissue-specific pathways contained genes that were associated with tissue-specific disease-risk SNPs in their 3’UTR. This highlights the strength of using PUMA networks to identify disease-related genes.

Another strength of PUMA is that it does not use correlations between miRNA expression levels and their target genes to model gene regulation. One of the reasons for not implementing correlation between regulators and their targets as an input in PUMA is that we have previously observed that a regulator’s expression level is often not associated with its regulatory potential [15], possibly due to combinatorial regulation of the target genes by multiple factors. The analysis presented in the current study again strengthens this. We believe that, while a miRNA needs to be expressed in order to regulate a target gene, the regulatory patterns of a miRNA are complex, and depend not only on the miRNA’s expression level itself, but on the entire collection of miRNAs that are available in a cell [40], as well as on the complete set of target mRNA transcripts that are expressed.

A good strategy to integrate PUMA networks with miRNA expression data is to overlay the network nodes with miRNA and target gene mRNA expression levels *after* the edges have been estimated with PUMA. This way, one would first identify tissue- or disease-specific edges, and then assess whether these are connected to highly or differentially expressed miRNAs. In fact, we recently used a similar approach to identify tumor suppressor genes downregulated by a cluster of non-coding elements, which had been associated with patient outcome in osteosarcoma [41].

Gene regulation is a complex process involving multiple factors, including both transcription factors and miRNAs. Understanding these regulatory processes, and how they change between phenotypes, helps elucidating the network changes that occur between health and disease. Identifying genes that are differentially regulated, but not necessarily differentially expressed, can help us to understand the likely potential that a given biological state has to respond to changes, including drug treatment or disease progression. Although there have been many attempts to model gene regulation by transcription factors, few methods have tackled miRNA regulation or both regulators together.

PUMA models gene regulation by miRNAs and transcription factors in a principled way by incorporating our understanding of the regulatory processes that control gene transcript levels. In applying PUMA to a wide variety of tissues, we find patterns of miRNA regulation associated with a variety of tissue-specific processes in ways that add explanatory power to the analysis of the same tissues using transcription factor regulation alone [15]. As such, PUMA provides the first robust computational method for modeling complex patterns of regulation involving miRNAs. Its implementation in freely-available, open-source code means that it can be broadly applied to the analysis of other phenotypes and disease states.

## ACKNOWLEDGMENTS

We would like to thank all members of the Quackenbush laboratory helpful discussions. M.L.K. was supported by a Charles A. King Trust Postdoctoral Research Fellowship Program, Sara Elisabeth O’Brien Trust, Bank of America, N.A., co-Trustees and by grants from the Norwegian Research Council, Helse Sør-Øst, and the University of Oslo through the Centre for Molecular Medicine Norway (NCMM). M.F. was supported by the project Investement for the future AMAIZING ANR-10-BTBR-01 (ANR-PIA AMAIZING). J.Q. was supported by a grant from the US National Cancer Institute (R35CA220523). K.G. was supported by a grant from the National Institutes of Health (K25HL133599).

## AUTHOR CONTRIBUTIONS

Conceptualization, M.L.K, J.Q., K.G.; Methodology, M.L.K., K.G.; Software, M.L.K., A.M., K.G.; Formal Analysis, M.L.K.; Investigation, M.L.K., M.F., K.G.; Resources, M.L.K., J.Q.; Data Curation, M.L.K.; Writing– Original Draft, M.L.K.; Writing–Review & Editing, M.L.K., M.F., A.M., J.Q., K.G.; Visualization, M.L.K., K.G.; Supervision, J.Q.; Funding Acquisition, M.L.K., J.Q, K.G.

---

**Supplemental Figure S1.**
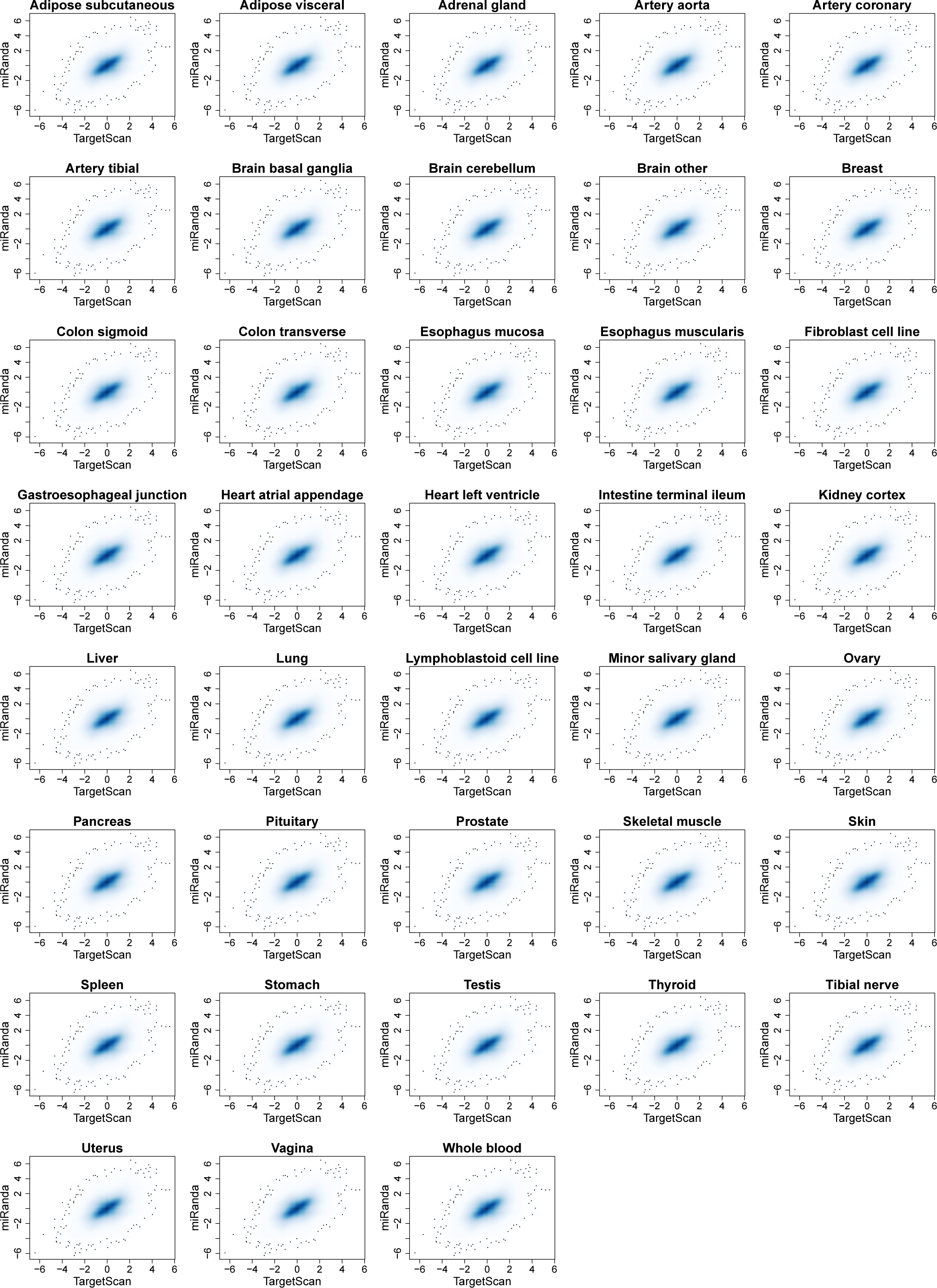
Smooth scatterplot depicting, for each tissue, the correlation of all tissue-specificity scores of networks modeled on the TargetScan and the miRanda prior.

**Supplemental Figure S2.**
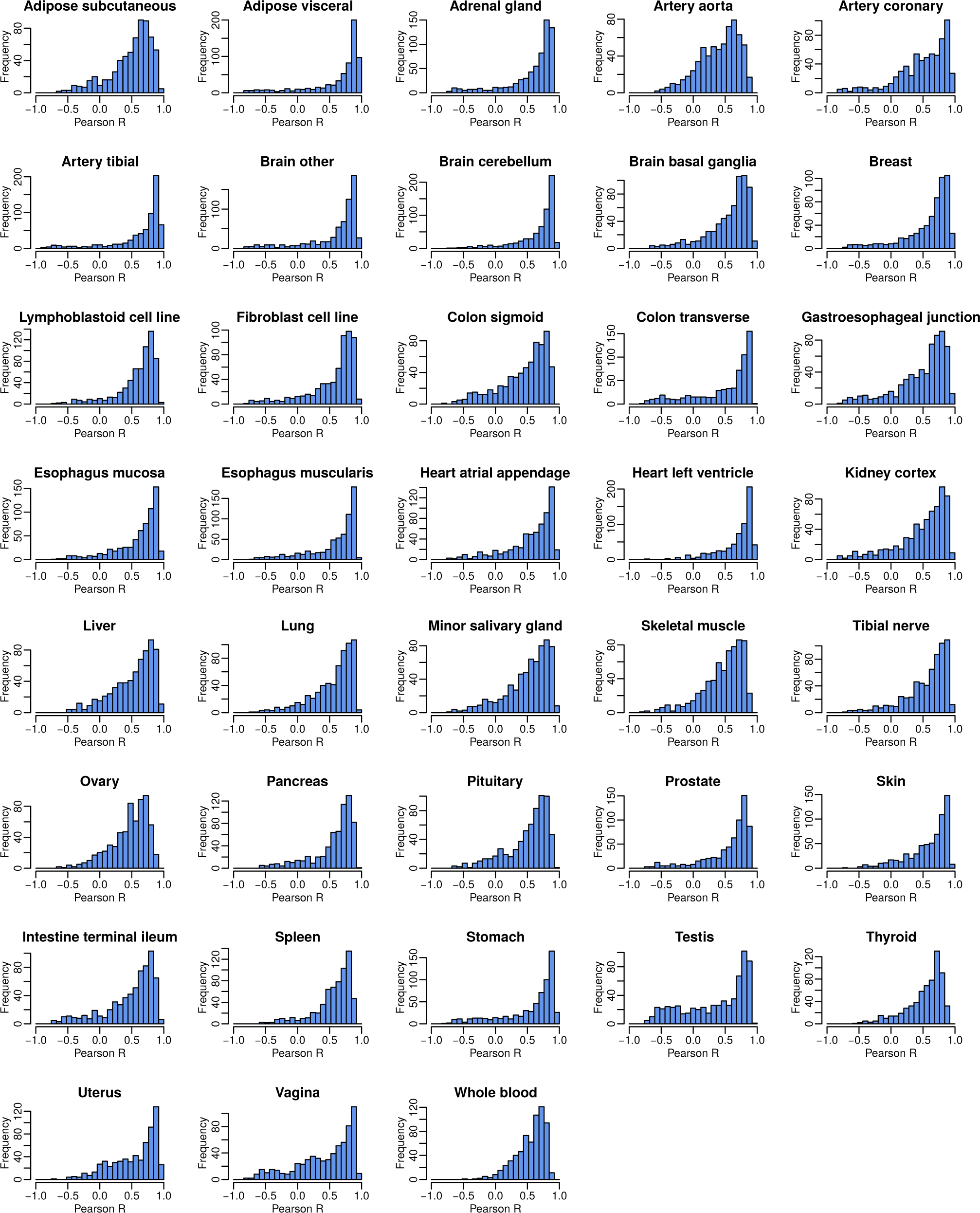
Histogram of Pearson correlation coefficients obtained from comparing the GSEA scores of the tissue-specific miRNA targeting profiles computed on the TargetScan and the miRanda prior, visualized for each tissue individually.

**Supplemental Figure S3.**
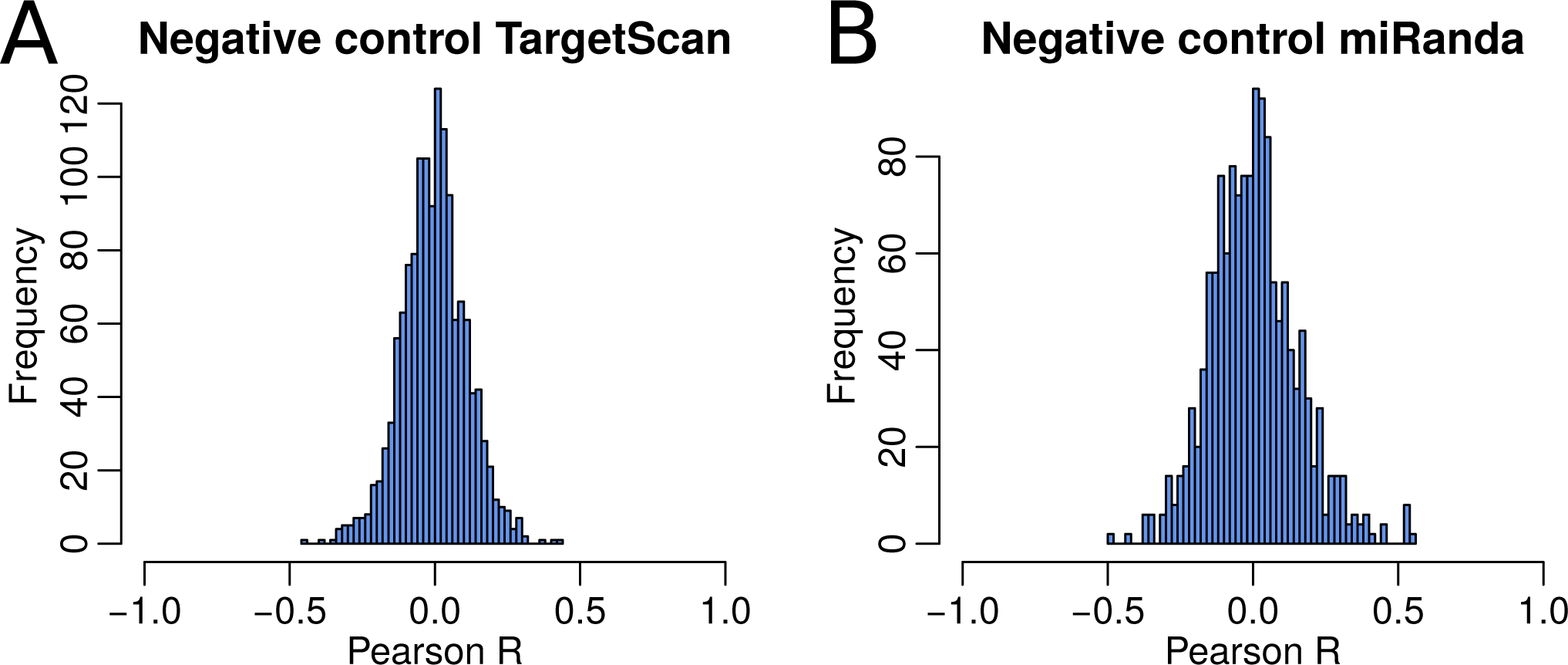
Negative control for the similarity analysis of miRNA/tissue GSEA scores predicted on networks obtained from the two different priors (shown in Figure 3). Here, we compared tissue-specific GSEA scores for one miRNA in one specific tissue with those from the same miRNA in all other tissues, using networks modeled on the same prior—either from TargetScan (A) or miRanda (B).

**Supplemental Table S1.**
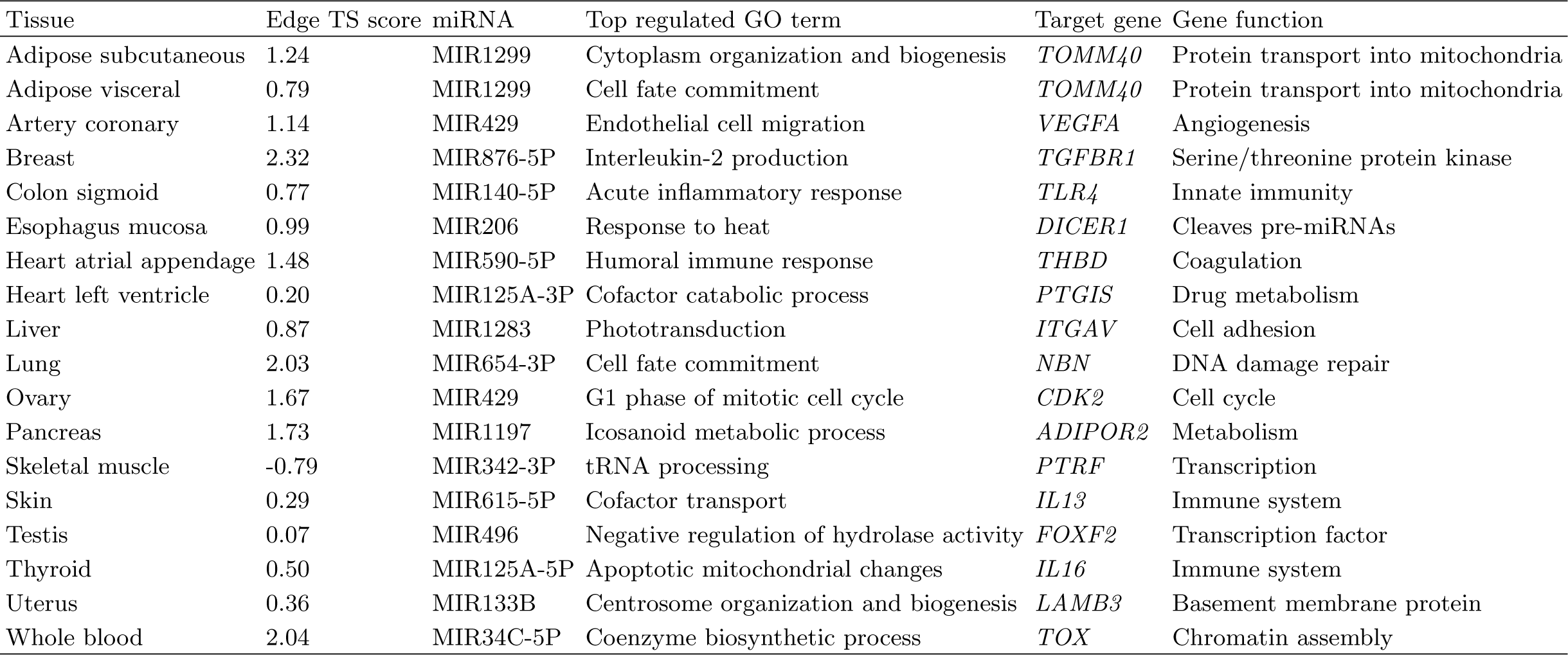
Top results from validation analysis using miRdSNP.

